# Compositional data analysis of microbiome and any-omics datasets: a revalidation of the additive logratio transformation

**DOI:** 10.1101/2021.05.15.444300

**Authors:** Michael Greenacre, Marina Martínez-Álvaro, Agustín Blasco

## Abstract

Microbiome and omics datasets are, by their intrinsic biological nature, of high dimensionality, characterized by counts of large numbers of components (microbial genes, operational taxonomic units, RNA transcripts, etc…). These data are generally regarded as compositional since the total number of counts identified within a sample are irrelevant. The central concept in compositional data analysis is the logratio transformation, the simplest being the additive logratios with respect to a fixed reference component. A full set of additive logratios is not isometric in the sense of reproducing the geometry of all pairwise logratios exactly, but their lack of isometry can be measured by the Procrustes correlation. The reference component can be chosen to maximize the Procrustes correlation between the additive logratio geometry and the exact logratio geometry, and for high-dimensional data there are many potential references. As a secondary criterion, minimizing the variance of the reference component’s log-transformed relative abundance values makes the subsequent interpretation of the logratios even easier. Finally, it is preferable that the reference component not be a rare component but well populated, and substantive biological reasons might also guide the choice if several reference candidates are identified. Results: On each of three high-dimensional datasets the additive logratio transformation was performed, using references that were identified according to the abovementioned criteria. For each dataset the compositional data structure was successfully reproduced, that is the additive logratios were very close to being isometric. The Procrustes correlations achieved for these datasets were 0.9991, 0.9977 and 0.9997, respectively. In the third case, where the objective was to distinguish between three groups of samples, the approximation was made to the restricted logratio space of the between-group variance. Conclusions: We show that for high-dimensional compositional data additive logratios can provide a valid choice as transformed variables that are (1) subcompositionally coherent, (2) explaining 100% of the total logratio variance and (3) coming measurably very close to being isometric, that is approximating almost perfectly the exact logratio geometry. The interpretation of additive logratios is simple and, when the variance of the log-transformed reference is very low, it is made even simpler since each additive logratio can be identified with a corresponding compositional component.

## Introduction

The article (1) is emphatically titled: “Microbiome datasets are compositional: and this is not optional”. We agree. For example, the number of so-called reads obtained by high throughput sequencing varies from sample to sample and is of no relevance to the investigation, much the same as the size of a rock is irrelevant to the study of its geochemical composition. It is the relative values of the read counts that are the data of interest, thus making the data strictly compositional (2). The same is true for other assay methods such as liquid chromatography–mass spectrometry where identification of metabolites is achieved by intensity values or areas under peaks.

It is convenient to eliminate the effect of the sample totals by normalizing, or *closing*, the data, so that sample values sum to 1 — these vectors of non-negative sample values with constant sums are called *compositions*. Once this initial step is made, the question remains how to analyze, relate and interpret the *components* of the compositions, be they microbial genes, operational taxonomic units, transcripts or metabolites.

It has long been appreciated, since the pioneering work of John Aitchison (3–5), that a valid, *(subcompositionally) coherent* way to tackle compositional data is by considering pairwise ratios of the components and by analyzing these ratios after logarithmic transformation. Notice that these ratios are invariant with respect to the normalization (closure) of the data. Coherence means that, if the set of components is extended or reduced, the ratios of the common components remain constant whereas the values of their relative abundances do change. In fact, the set of components under consideration, imposed by the measuring instrument, research objective and practical considerations, is almost always a subcomposition of a potentially much larger set.

The basic concept and data transformation in compositional data analysis is thus the *logratio*, the logarithm of pairwise ratios, with the log-transformation serving to change the ratio scale to an interval one: log(*A/B*) = log(*A*) *−* log(*B*). The challenge is to choose a data transformation that replaces the compositional dataset with a set of logratios that are substantively meaningful to the practitioner as well as having a clear interpretation. Once the transformation to logratios is performed, analysis, visualization and inference carries on as before, but always taking into account the interpretation in terms of ratios.

In Aitchison’s earliest work he proposed the additive logratio transformation (ALR), where one component is chosen as the denominator, or *reference*, with all the other components as numerators. Thus, if there are *J* components, with values *X*_1_, *X*_2_, …, *X*_*J*_, there are *J −* 1 logratios in the ALR set with respect to the selected reference component, denoted by *ref*, of the form:

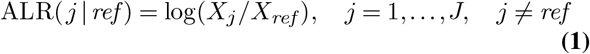

Since then a variety of logratio transformations have been proposed: for example, centered logratios (used in (6)), isometric logratios and pivot logratios (for example, (7, 8)). All of these involve ratios of geometric means of components and, as a result, have complicated interpretations (9, 10), lacking the simplicity of the pairwise logratio between two components. Isometric and pivot logratios are particularly problematic when the numbers of components in the geometric means are high. They do have the property of isometry, however, which means that they engender exactly the same multivariate geometric structure of the sample points as that of all the pairwise logratios, called the *logratio geometry* (sometimes referred to as the “Aitchison geometry”). The proponents of these complex transformations take isometry as a type of “gold standard” for the analysis of compositional data, and the strict adherence to this mathematical ideal has been to the detriment of using simpler transformations such as the ALRs, or a subset of pairwise logratios. In a series of papers (11–13) it is shown in a variety of contexts that a set of simple pairwise logratios can satisfactorily approximate the logratio geometry, coming close to being isometric for all practical purposes. A small loss of isometry is thus traded off in favour of the benefit of a simpler and clearer interpretation of the logratio variables. In these above-mentioned studies any set of pairwise logratios can be selected, whereas ALRs are restricted to pairwise logratios with respect to a fixed reference component.

Apart from the fact that ALRs are not strictly isometric, various other criticisms have been levelled at the ALR transformation, such as its sacrificing a component to serve as the reference and the doubt about which component to choose as reference. We hope to show that none of the above are disadvantages, but rather that, especially in the case of high-dimensional compositional data, the ALRs are the logratio transformations of choice and that their involving a fixed reference is actually a benefit. In this way we return to the origins of compositional data analysis and re-establish the additive logratio in all fields of omics research, thereby vindicating Aitchison’s original claim as enounced in the following quotation from his keynote address (14) at the biennial Compositional Data Analysis workshop in 2008 (Section 5.1):

> “The ALR transformation methodology has, in my view, withstood all attacks on its validity as a statistical modelling tool. Indeed, it is an approach to practical compositional data analysis which I recommend particularly for non-mathematicians. The advantage of its logratios involving only two components, in contrast to CLR and ILR (isometric transformations …), which use logratios involving more than two and often many components, makes for simple interpretation and far outweighs any criticism, more imagined than real, that the transformation is not isometric.”

Aitchison’s phrasing above that the criticism of the ALR transformation not being isometric is “more imagined than real”, is particularly pertinent to what we will show here. We will demonstrate that a set of ALRs can be so close to being isometric that, for all practical purposes, they are isometric. We will also show that there are clearly defined criteria for choosing a reference and it is advantageous that there are very many potential choices in high-dimensional data when the number of components is large.

Three high-dimensional omics datasets will be used to show that the ALR transformation can validly provide a set of simple variables to represent the whole compositional dataset, the essential step being the choice of the reference component. Section 2 gives some background theoretical material. Section 3 details the computational steps involved in determining and validating the ALRs chosen for each of three datasets, and Section 4 gives the results. Section 5 closes with a discussion and conclusion.

## Logratio variance, logratio geometry and selection of additive logratio transformation

Logratio-based compositional data analysis, often called CoDA (7), has mainly developed in fields where the number of components *J* is less, often much less, than the number of samples *I*, i.e. *J < I*, with geochemistry being the area of most applications. A short, yet comprehensive, review of CoDA is given by (15), with recent books aimed at practitioners by (16) and (8). The relevant theoretical results for our purpose are summarized in this section, as well as how they apply to ALRs.

### Total logratio variance

The total logratio variance is a basic statistic that quantifies how dispersed the samples are in the multivariate logratio space. A compositional data vector with *J* components, *X*_1_, *X*_2_, …, *X*_*J*_, can be expanded into 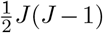 pairwise ratios, and then log-transformed. Thus, an *I* × *J* compositional data matrix can be expanded, notionally at least, to an 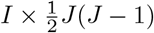 matrix of logratios. In the most general case, there are positive weights *c*_1_, *c*_2_, …, *c*_*J*_ associated with the components (17–20), where *c*_1_ + *c*_2_ +… + *c*_*J*_ = 1, in which case it can be shown that the (*j, k*)-th logratio log(*X*_*j*_*/X*_*k*_) has weight equal to the product *c*_*j*_*c*_*k*_ (15, 16). The total logratio variance is then defined as:

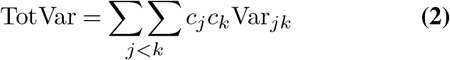

where Var_*jk*_ is the variance of the (*j, k*)-th logratio (15, 16). The weights have a normalizing function to balance out the contributions of the different components, since rarer components often engender excessively large logratio variances (2, 16, 21), or they might be used to downweight components with high measurement error. However, in many applications, including the ones in this article, this aspect is ignored and the components are equally weighted by *c*_*j*_ = 1*/J, j* = 1, …, *J*. Consequently, (2) simplifies as the sum of the 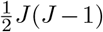 variances of the unique pairwise logratios divided by *J* ^2^.

For a dataset with thousands of components this would be a laborious calculation, but fortunately there is a shortcut thanks to the centred logratio (CLR) transformation:

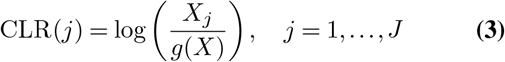

where *g*(*X*) is the weighted geometric mean 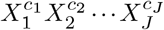, (16) i.e.

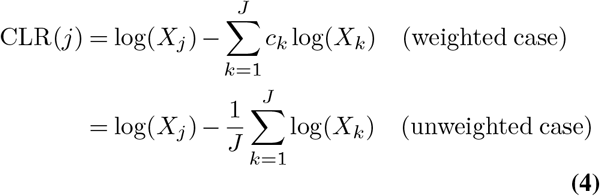

The total variance in (2) is then equivalently computed using the variances of the CLRs, Var_*j*_, weighted respectively by *c*_*j*_, *j* = 1, …, *J*, or by constant 1*/J* when equally weighted:

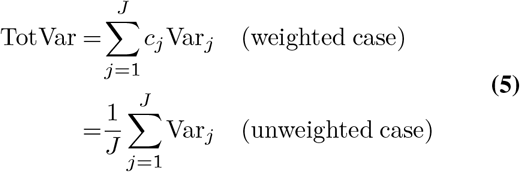

Notice that in the weighted or unweighted cases the CLRs have to be computed according to one of the respective definitions in (4). Notice too that the weighting in (2) or (5), with either differential or equal weights, are weighted averages of the part variances, ensuring that total logratio variances can be compared between data sets of different sizes.

The computation is completely symmetric with respect to rows and columns, so when *J > I*, as will generally be the case for omics data, the computation can be further simplified. The data matrix is first transposed and relative abundances are expressed with respect to component totals, following which the above computation is repeated as if the samples were the components.

### Logratio geometry

A compositional dataset has a certain exact geometry defined by the logratio distances between every pair of samples. These are Euclidean distances that can be defined in two equivalent ways: either on the 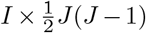 matrix of all pairwise logratios, again a very wide matrix due to the large number of pairs of components, or more efficiently on the *I* ×*J* matrix of CLRs (4). As before, there are weighted and unweighted versions — for the exact definitions see (15, 16)). If *J* < *I* (i.e. the dataset is “narrow”) the sample points are exactly in a (*J* −1)-dimensional Euclidean space, otherwise if *J* > *I* (i.e. the dataset is “wide”) they are exactly contained in a (*I*− 1)-dimensional Euclidean space — hence, the dimensionality is *K* = min {*I*− 1, *J* −1}.

In both weighted and unweighted cases the total logratio variance can be decomposed along principal axes to give a low-dimensional reduced view of the samples, called *logratio analysis* (LRA) (22). LRA is the principal component analysis of all the pairwise logratios, which is equivalent to the PCA of all the CLRs, in weighted (20) or unweighted (23) forms.

Notice that for a compositional data set of dimensionality *J* −1, say (for the case *J* <= *I*), then any set of *J*− 1 linearly independent logratios, including any set of *J*− 1 ALRs, explains the total logratio variance in 2 or 5 completely. This set clearly does not contain the total variance, but explains it totally in a regression sense (11). If *J* > *I*, as in many high-dimensional datasets, only *I* − 1 linearly independent logratios are required to explain the total logratio variance.

### Procrustes analysis

For any particular set of logratio transformations, the samples in the transformed space can be “fitted” to the exact logratio geometry, using *Procrustes analysis* (24, 25), to see how close they come to the exact geometry. Suppose the coordinates of the samples in their exact logratio geometry are in the matrix **X** (*I* × *K*), where *K* is the dimensionality of the space, as explained above. The coordinates are established using LRA and the inter-sample distances in this geometry are exactly the logratio distances. Similarly, suppose the coordinates of the samples in a particular ALR geometry are in the matrix **X** (*I* × *K*), the same dimensionality as the exact one — for example, if *J* > *I* (as in the present case) then the dimensionality of the logratio space is *K* = *I* − 1 (one less than the number of samples), and that of the *J* − 1 ALRs, also involving *I* samples, is also *I* −1. The sample coordinates in the ALR geometry are established using PCA and the inter-sample distances in this ALR geometry will not be the same as the exact logratio distances, partly due to differences in scale and rotation between the two matrices, which are irrelevant to summarizing their distance structure. So Procrustes analysis aims to match the configurations by least-squares as closely as possible by three simple operations: centering, scaling and rotation.

The first two operations are trivial: the columns of **X** and **Y** are already centered by the LRA and PCA respectively, and scaling is achieved by dividing each matrix by the square roots of their respective sum-of-squares. Suppose **X**^*^ and **Y**^*^ are the matrices standardized in this way, then compute the singular value decomposition of their cross-product (**X**^*^)^T^**Y**^*^ = **UDV**^T^. The fitting of **Y**^*^ to **X**^*^ by leastsquares fitting is achieved by applying the rotation matrix **Q** = **VU**^T^ to **Y**^*^: **Y**^*^**Q**. Equivalently, **X**^*^ could be fitted to **Y**^*^ by applying the inverse rotation **Q**^T^ : **X**^*^**Q**^T^.

The final step is to compute the Procrustes correlation, which measures how close the two configurations are to being exactly matched. The sum-of-squares *E* of the differences between **X**^*^ and **Y**^*^**Q** lies between 0 and 1, where 0 implies perfect matching and 1 implies total absence of matching. The quantity *E* can be considered a residual sum-of-squares if one thinks of **Y**^*^ being fitted to **X**^*^, and since *E* has a maximum of 1, then 1 − *E* is analogous to a coefficient of determination (*R*^2^) in a least-squares regression. The Procrustes correlation is thus defined as 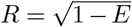, so that a value near 1 would mean that the ALR geometry is very close to the exact logratio geometry, that is it is almost *isometric*. The Procrustes correlation *R* can be equivalently computed as the correlation between the elements of the matrices **X**^*^ and **Y**^*^**Q** strung out as *IK* × 1 vectors.

In short, the goal is to measure the deviation of the ALR-transformed data from the ideal of isometry. This way of measuring the proximity between two configurations in multidimensional space by the Procrustes correlation has already been used to select a subset of pairwise logratios that engenders a Euclidean geometry close to the exact one (11–13). This idea was inspired by the selection of variables in PCA (26), and the same idea will be used here to select a reference in order to define a set of ALRs.

### Criteria for selecting reference for additive logratios

The ALR transformation converts the original *I* × *J* compositional data matrix to an *I* × (*J−* 1) matix of ALRs, with respect to a particular reference component. There are *J* potential reference components to choose from, which in the usual geo- and biochemical applications can be a relatively low number. However, in the case of most omics data, *J* is very large and usually very much larger than *I*, the number of samples. This gives a large set of possibilities for choosing a set of ALRs that comes as close as possible to reproducing the exact logratio geometry by achieving a very high Procrustes correlation.

The matching of the geometries is the most important criterion for choosing the reference, but there are other properties that would be desirable.. For example, it would be very convenient if the reference’s relative abundances across the samples is as constant as possible. From (1) ALR(*j* |*ref*) = log(*X*_*j*_) *−*log(*X*_*ref*_), hence we should look for low variance in log(*X*_*ref*_). Since dividing each component by an almost constant reference value just shifts all the logratios by an almost constant amount, the logratio can then be interpreted in practice as its numerator on a logarithmic scale. An additional benefit of choosing a low variance component is that it is unlikely to be correlated with any continuous or categorical covariate whose relationship with the compositions is being investigated — the actual relationship with such covariates can be checked where applicable.

A further criterion would be to avoid choosing a reference with low occupancy across individuals, where low occupancy is related to low overall abundance (27). Zeros need to be replaced before making the logratio transformation, by adding one to all the counts or using one of the many zero replacement methods, and using such a component as the denominator would affect the interpretation of all the ALRs.

## Validating the ALR transformation on three datasets

Three datasets with high numbers of components are considered here:

- a wide functional microbe dataset of secum samples of *I* = 89 rabbits, in a study of *J* = 3937 microbial genes (28), which we will refer to as the Rabbits data;
- a wide dataset of *I* = 28 people in a study of *J* = 3147 mRNA transcripts from mouse bone marrow dendritic cells (29), re-analysed by (21), which we will refer to as the RNAseq data;
- a narrower dataset, from the stool samples of *I* = 490 patients in a study of colono-rectal cancer, on which *J* = 335 microbiota operational taxonomic units (OTUs) were identified, referred to as the Cancer data (30); the individuals in this dataset were classified as normal, adenoma (benign tumor) or cancer and the objective was to identify OTUs that discriminated between these groups, especially to predict the cancer group.

The third dataset is included because it involves much fewer components than the others and problematically has more than 50% data zeros. It also allows investigating how well ALRs can reproduce the geometry of the subspace of the between-group variance, since variance unrelated to the group discrimination is not of interest in the study.

For each dataset the following statistics are computed:

a. The total logratio variance, which is a statistic that summarizes how dispersed the sample points are in multidimensional space (equal weighting of components will be used throughout). For the first two wide examples, the total variance can be more efficiently computed by transposing the matrix of abundances (or relative abundances) and then computing the total variance on the CLRs of the samples, as if they were the components. The exact logratio geometric structure is then determined, that is the coordinates of all the sample points in the full space of all the dimensions (for first two datasets), or in the constrained space (third dataset). And then, for each component used as a reference for defining ALRs:
b. The Procrustes correlations between the exact logratio geometry and the approximate geometry of the set of ALRs engendered by each choice of reference.
c. The variance of the log-transformed relative abundances of each reference candidate across the samples;

The components with the highest correlations in (b) and, of those, the lowest variances in (c) will be candidates for the choice of reference. In practice, of course, domain knowledge should also play a role in selecting the reference, especially when there are several competing candidates.

Finally, we will show the reduced-dimension LRA of the exact sample configuration based on all pairwise logratios alongside the reduced-dimension configuration of the chosen set of ALRs to demonstrate that the configurations are practically identical.

## Results

### The Rabbits data

This is a 89 × 3937 dataset of counts and there are no zeros.

a. Total logratio variance = 0.1601, computed on the 3937 CLRs of the components (microbial genes). Equivalently, a faster way is to transpose the dataset and then treat the samples as components — the same result is obtained on the 89 CLRs of the samples.
b. The highest Procrustes correlation is equal to 0.9991, corresponding to the same gene number 856. This gene has the 201st highest abundance among the 3937 genes.
c. The lowest variance of the log-transformed relative abundance of the reference components is equal to 0.00117, corresponding to gene number 856. Its five-point summary on the log-scale is:

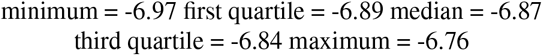

showing a high constancy in the values, with interquartile range of 0.05.

To visualize how close the ALR variables are being to isometric, Figure 1 shows the all between-sample distances computed on the ALRs plottted against the exact logratio distances based on either all pairwise logratios or, equivalently, the CLRs.

**Fig. 1.**
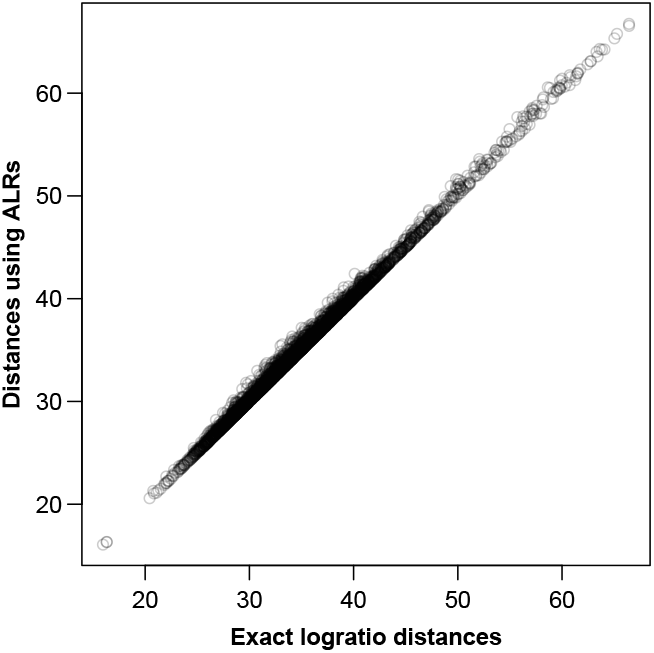
Between-sample distances for the Rabbits dataset based on the ALRs with reference microbial gene 856 versus the exact logratio distances, corresponding to the Procrustes correlation of 0.9991. The number of pairs of distances plotted = 89 × 88*/*2 = 3916.

The LRA of the full dataset, showing just the samples, is shown in Figure 2a, while the corresponding PCA of the ALRs with reference gene 856 is shown in Figure 2b. They are practically identical, with very slight differences, as expected. The letters S and F stand for the two laboratories that did the sequencing, showing a clear separation. This sequencer effect was subsequently eliminated in the data analysis.

**Fig. 2.**
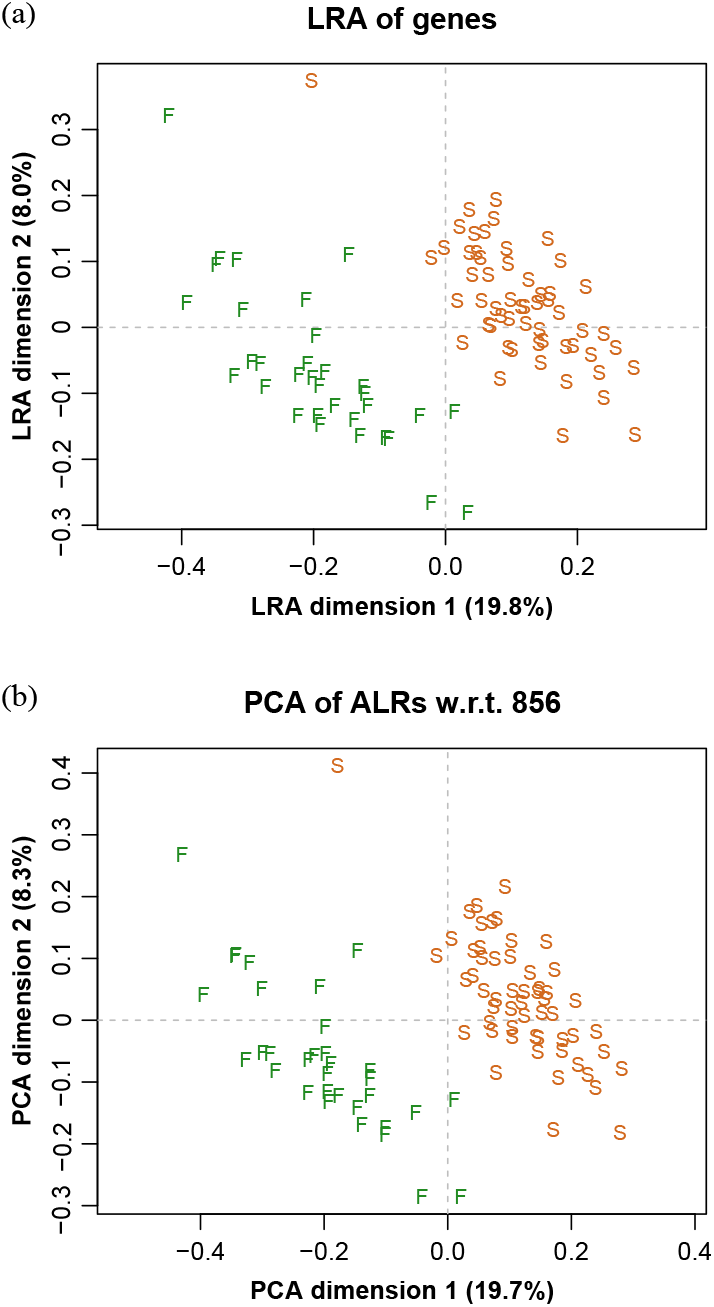
(a) Logratio analysis of the Rabbits data, aiming to explain the total logratio variance. (b) Principal component analysis of the additive logratios with reference component microbial gene number 856, showing a geometry practically identical to the exact logratio geometry. The two groups of points are due to two sequencing laboratories, indicated here by F and S.

The low variance of the reference gene means that in the original table of counts this gene’s counts are closely proportional to the total counts — Figure 3 shows this conclusively.

**Fig. 3.**
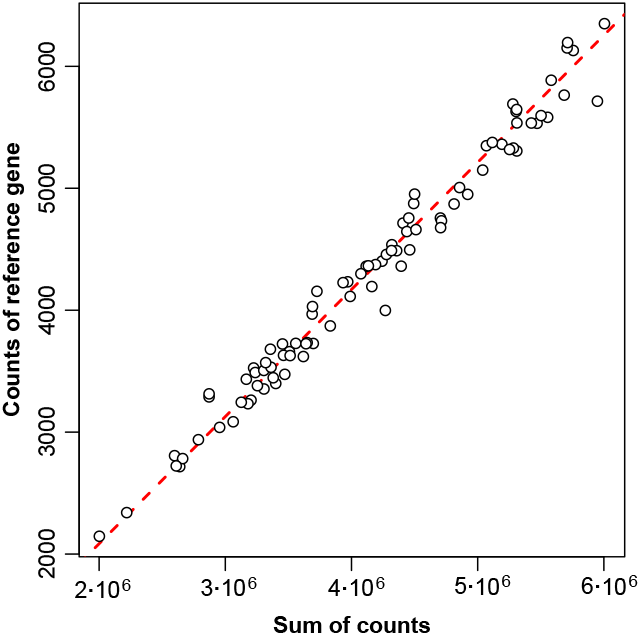
Proportionality between counts of reference gene number 856 and sum of counts, for the 89 samples. The dashed red line goes through the origin (0,0).

It is interesting that the top candidates in this data set coming up as reference microbial genes are associated with the genetic machinery of the microbes (28), which are intrinsic in all microbial ecosystems. The same pattern has been found for other functional microbiome datasets (31).

### The RNAseq data

This is a 28 ×3147 dataset of counts. There are 32 zeros in this dataset, which have been replaced using the function cmultRepl in R package zCompositions (32).

a. Total logratio variance = 0.2099, computed on the 3147 CLRs of the components (transcripts). Equivalently, a faster way is to transpose the dataset and then, after treating the samples as components, the same result is obtained on the 28 CLRs of the samples.
b. The highest Procrustes correlation is equal to 0.9977, corresponding to the mRNA transcript number 1318.
c. The lowest variance of the log-transformed relative abundance of the reference components is equal to 0.00415, corresponding to mRNA transcript number 1557. Its five-point summary on the log-scale is

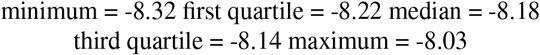

showing again a high constancy in the values, with interquartile range of 0.08.

In this case the reference that maximizes the correlation is different from the one that minimizes the variance. One transcript, number 1179, comes second on both criteria and is the one that was chosen, with Procrustes correlation = 0.997 and variance = 0.00626. It has the 1617th highest relative abundance among the 3147 transcripts, and its five-point summary is:

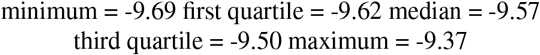

with interquartile range 0.12.

To visualize how close the ALR variables are being to isometric, Figure 4 shows the between-sample distances computed on the ALRs plotted against the exact logratio distances based on either all pairwise logratios or, equivalently, the CLRs. The agreement is again excellent, with slightly less congruence in the high distances (commented below).

**Fig. 4.**
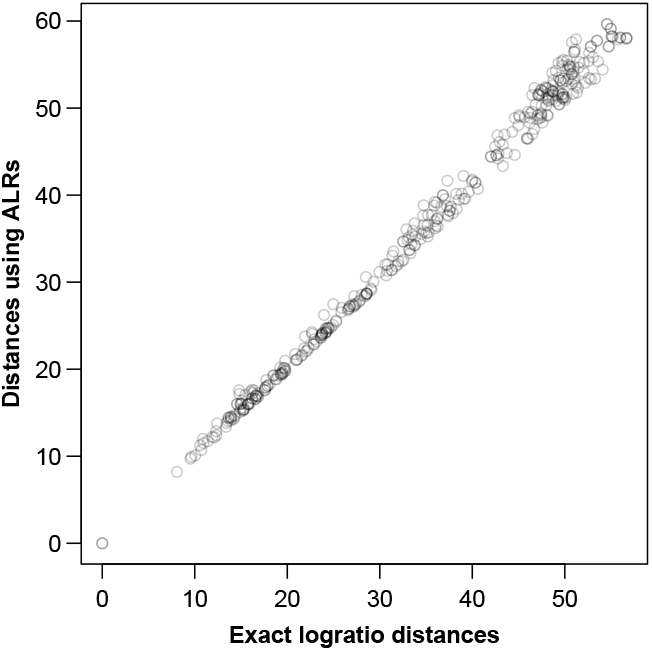
Between-sample distances for the RNAseq data based on the ALRs with reference transcript 1179 versus the exact logratio distances, corresponding to the Procrustes correlation of 0.9977. The number of pairs of distances plotted = 28 27/2 = 378.

The LRA of the full dataset, showing just the samples, is shown in Figure 5a, while the PCA of the ALRs with reference transcript 1179 is shown in Figure 5b. They are practically identical, with only very slight differences, again as expected from the very high Procrustes correlation. The labels stand for two different treatments (L and M) and 7 different times (0, 1, 2, 4, 6, 9 and 12 hours). The slight discrepancies in the higher distances of Figure 4 would correspond to the distances between samples of the different treatment groups, which are the most separated in Figure 5.

**Fig. 5.**
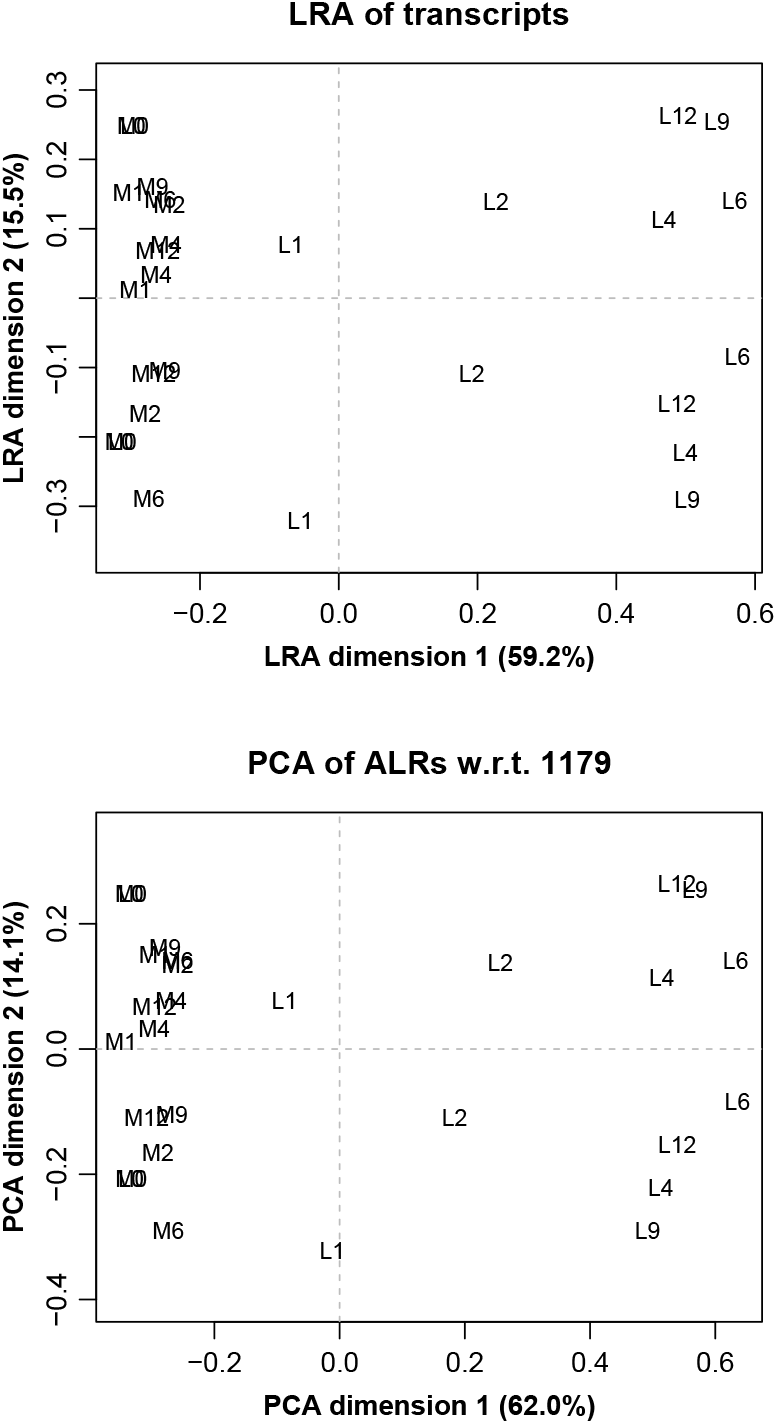
(a) Logratio analysis of the RNAseq data, aiming to explain the total logratio variance. (b) Principal component analysis of the additive logratios with reference transcript number 1179, which has a geometry almost identical to the exact logratio geometry. The label prefixes M and L refer to two treatments, and the number suffixes refer to times in hours.

### The Cancer data

This is a 490 × 335 dataset of counts. This dataset has a large number of zero counts (58% of the whole dataset) — a one was added to all the counts in the dataset before closing and taking logarithms. One OTU was removed due to a very large outlying value, which might be incorrect, reducing the dataset to 334 OTUs. The samples were divided into three groups: adenoma (benign tumor), cancer and normal.

This is a more problematic example because of the fewer components and the presence of so many zeros. In a preliminary analysis, the best Procrustes correlation obtained with the full-space geometry (of dimension 333, one less than the number of ALRs) was 0.936, lower than those of the first two examples. Since the objective of the data is to find the OTUs that discriminate between the groups, agreement in the geometry is not required in the full space but rather in the two-dimensional reduced space of the group means, excluding the dimensions not related to group differences. So, in the search for maximum agreement, a constrained (or restricted) LRA is performed on the complete set of CLRs, constrained to the three group means, and compared with the series of similarly constrained PCAs of ALR-transformed data using the different reference components. In all cases we are not interested here in variance that it is not related to the separation of the three sample groups.

a. Total logratio variance = 1.5301 computed on the 334 CLRs of the components, which in this example are less than the number of samples. Notice that this value is much higher than that of the previous two datasets, which is typical of taxa datasets, with many zeros and larger contrasts in the data. However, the restricted between-group variance that we focus on here is small, equal to 0.0125, only 0.82% of the total. Nevertheless, the group differences are highly significant (*p <* 0.0001), using the multivariate permutation test in the vegan package (33).
b. The highest Procrustes correlation is equal to 0.9997, corresponding to the OTU number 312 (labelled in the original dataset as Otu000363).
c. The lowest variance of the log-transformed reference components is equal to 0.308, corresponding to OTU number 320 (labelled in the original dataset as Otu000372). Its five-point summary on the log-scale is

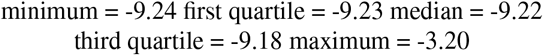

Since this OTU has only 36 nonzero values out of 490, the above estimated quartiles relate mostly to the overwhelming number of zeros, hence the small differences in quartiles up to the third, with positive skewness up to the maximum corresponding to the nonzero values.

It was decided to use this OTU number 312 as the reference part, with variance of its log-transformed relative abundances equal to 0.476 and many more nonzero abundances, 213 out of 490. Its five-number summary is as follows:

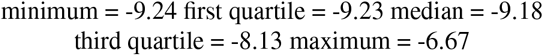

Because of the variability in the reference part, the ALRs should always be interpreted as pairwise logratios with respect to the reference, not as approximating the logarithms of the numerator components as in the first two examples.

The Procrustes correlation almost equal to 1 again means that the ALRs are, for all practical purposes, isometric, in this case isometric for group discrimination.

Figure 6 shows the constrained solution of the between-group variance using all the pairwise logratios (i.e. the centred logratios), and the corresponding solution using the ALRs. Because of the Procrustes correlation close to 1, there is no noticeable difference between the two solutions. The fact that the confidence ellipses for the groups means are highly separated bears testimony to the highly significant differences between them. For each axis two percentages are given: the first is the explained variance relative to the two-dimensional constrained logratio variance (= 0.0125), the second is the same explained variance relative to the much larger total logratio variance (= 1.5301).

**Fig. 6.**
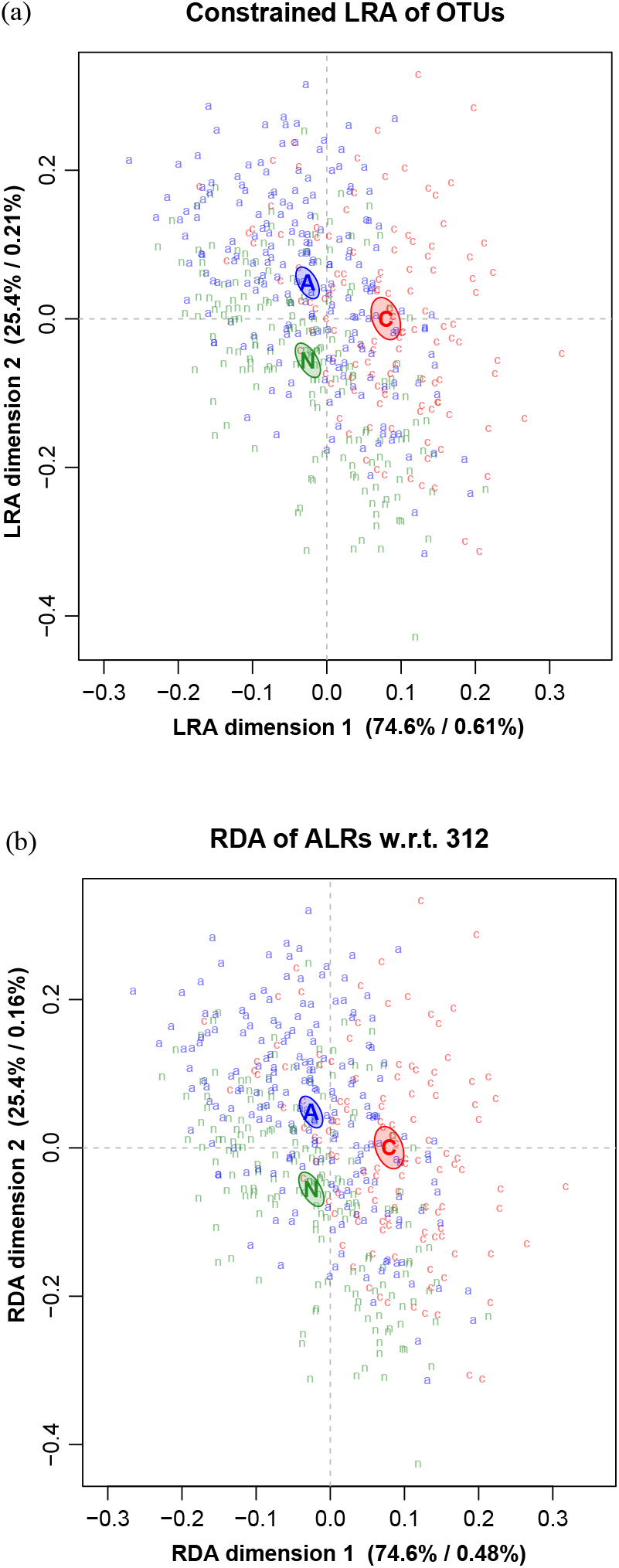
(a) Constrained logratio analysis of the Baxter data, aiming to explain the between-groups logratio variance. (b) Constrained principal component analysis (i.e. RDA) of the additive logratios with reference OTU number 312. In each case the first percentage expresses explained variance relative to the constrained logratio variance, whereas the second percetnage is relative to the total logratio variance.

In this example there were much fewer components to choose from, and there were a large number of data zeros, typical of a microbial taxa dataset. These data zeros force samples onto the sides of the simplex space of the original compositions, and in the logratio transformations these become outliers, hence the difficulty in matching the exact and approximate geometries in the full space of the dataset. When constraining the solution to discriminate between the three groups, however, the outliers are much less important and so the ALRs, with the reference that was chosen to produce a configuration close to the constrained solution, has functioned surprisingly well, as shown in Figure 6.

Having made a selection of a set of ALRs that reproduces the between-group geometry, these ALRs can be used in a model for predicting the groups. In (30) a random forest prediction algorithm is used, combined with a backward elimination of the components, using the R package AUCRF (34) to get an optimal subset of OTUs. The classification is aimed at predicting the cancer and normal groups (hence the ademona group is omitted). Using the relative abundances as inputs to the algorithm, this results in 33 OTUs being chosen, and an overall error rate of 22%, with 52% of the cancers correctly predicted, and 97% of the normals. As a comparison, using the ALRs with respect to reference 856, the same algorithm with the same decision rule results in 34 OTUs being chosen, and an overall error rate of 20%, with 66% of the cancers correctly predicted and 90% of the normals. Moreover, the list of chosen OTUs is very similar (i.e. in the case of the ALRs, the denominator OTUs), with the top 10 in each selection identical. At least in this example, using ALRs has performed slightly better, with the added value that logratios are being used, conforming to good practice in compositional data analysis.

The use of the relative abundances in (30) seems to be hardly different than the logratio approach, at least for the variable selection and prediction. Correspondence analysis (CA), which also operates on the relative abundances, has also been applied to this dataset (15) and shown to produce similar results. There is a theoretical reason underpinning CA, however, and that is the fact that the chi-square normallization in CA has a close connection to logratio distance, and the issue of lack of subcompositional coherence is less critical (35–37). The striking advantage of CA for compositional data analysis is the fact that it handles zero values naturally, which is in fact why it is so popular in ecology and other fields where data can be very sparse, such as archaeology and linguistics.

## Discussion

Our objective has been to show that the ALR transformation, the simplest one in the CoDA toolbox, can provide a valid solution for the analysis of high-dimensional compositional datasets. The challenge is to find a good reference part.

In the two datasets with more than 3000 components, there was more chance to find a reference component with the two desirable properties for constructing a set of ALRs: first and foremost, the reference has to result in a high Procrustes correlation between the exact logratio geometry and the ALR geometry, both of which have the same dimensionality. A secondary criterion is low variance in the log of the relative abundance, which considerably simplifies the interpretation of the ALRs. In all three examples we have found that the ALR transformation has proved to be suitable for representing the variability of the compositional dataset, either this variability in the full space of the data, or in a space that is constrained to the particular objective of the study.

There has been a rejection in the compositional data analysis literature of variables that are not exactly isometric, and variables that are “oblique” (10). This criticism is difficult to understand when one can come up with a set of variables that reproduces almost perfectly the logratio geometry, which means that the criticism is aimed at the tiny lack of isometry.

With respect to the ALRs, which are of concern here, these are simple pairwise logratios with respect to a chosen reference. If one is fortunate to find a reference that is almost constant in its relative abundance, this means that the pair- wise logratio in each ALR is, for all practical purposes of interpretation, the same as the logarithm of the numerator. This makes the interpretation of the ALRs much easier when it comes to judging which ALRs are important for explaining variance, relating to covariates or distinguishing between groups.

For the third example with hundreds of components and many zeros, it seems that the “best” reference (in the sense defined here) might not be as successful in reproducing the logratio geometry as perfectly as when there are thousands of components, especially when the reference itself contains many zeros. Thus, one has to carefully evaluate the properties of the ALRs as a potential set of logratios in any given study. For this particular dataset, however, with its principal objective of distinguishing between sample groups, the geometry of the space constrained to the variance between groups is very successfully reproduced by a set of ALRs. It appears that imposing constraints on the geometry according to the research objective, “regularizes” the space by removing the many problems of the zeros and consequent outliers.

We have shown that the ALR transformation can validly be used for high-dimensional datasets, and considerably simplify the life of practitioners. Hron et al. (10) state that “alr coordinates cannot be simply identified with the individual original components, as they are in fact logratios, but the link with these is more clearly stated”. We have shown that this sweeping statement is in fact not true in some cases. When the reference is almost constant, then the numerators of the ALRs are very close to being directly interpretable as the log-transformed relative abundances of the respective components. Then, for all practical purposes, the ALR can be referred to as the component itself. In addition, variances and correlations of the ALRs can be identified approximately with those of the numerator, apart from an overall scale factor, which makes the interpretation much easier. This has been possible for the first two datasets presented here, with the caveat that for the third set of microbial taxa counts, the reference part does not have sufficiently low variance for this simplified interpretation, and thus the ALRs should be interpreted as true ratios.

